# Contextualising samples: Supporting reference genomes of European biodiversity through sample and associated metadata collection

**DOI:** 10.1101/2023.06.28.546652

**Authors:** Astrid Böhne, Rosa Fernández, Jennifer A. Leonard, Ann M. McCartney, Seanna McTaggart, José Melo-Ferreira, Rita Monteiro, Rebekah A. Oomen, Olga Vinnere Pettersson, Torsten H. Struck

**Affiliations:** Leibniz Institute for the Analysis of Biodiversity Change, Museum Koenig Bonn, Centre for Molecular Biodiversity Research, Adenauerallee 127, 53113 Bonn. Germany; Metazoa Phylogenomics Lab, Biodiversity Program, Institute of Evolutionary Biology (IBE, CSIC-UPF), Passeig maritim de la Barceloneta 37-49, 08003, Barcelona, Spain.; Conservation and Evolutionary Genetics Group, Estación Biológica de Doñana (EBD-CSIC), Avda. Americo Vespucio 26, 41092, Sevilla, Spain.; Genomics Institute, University of California Santa Cruz, Santa Cruz, CA, USA; Earlham Institute, Norwich Research Park, Norwich, Norfolk, NR4 7UZ, United Kingdom; (1) CIBIO, Centro de Investigação em Biodiversidade e Recursos Genéticos, InBIO Laboratório Associado, Campus de Vairão, Universidade do Porto, 4485-661 Vairão, Portugal; (2) Departamento de Biologia, Faculdade de Ciências, Universidade do Porto, 4099-002 Porto, Portugal; (3) BIOPOLIS Program in Genomics, Biodiversity and Land Planning, CIBIO, Campus de Vairão, 4485-661 Vairão, Portugal.; Leibniz Institute for the Analysis of Biodiversity Change, Museum Koenig Bonn, Centre for Molecular Biodiversity Research, Adenauerallee 127, 53113 Bonn, Germany.; (1) Centre for Ecological & Evolutionary Synthesis, University of Oslo, Blindernveien 31, 0371 Oslo, Norway, (2) Natural History Museum, University of Oslo, P.O. Box 1172, Blindern, 0318 Oslo, Norway, (3) Centre for Coastal Research, University of Agder, Universitetsveien 25, 4630 Kristiansand, Norway, 4) Department of Biological Sciences University of New Brunswick Saint John, 100 Tucker Park Road E2K5E2, (5) Tjärnö Marine Laboratory, University of Gothenburg, Hättebäcksvägen 7 45296.; Science for Life Laboratory - Sweden (SciLifeLab), National Genomics Infrastructure, Uppsala University, P.O. Box 815, SE-752 37 Uppsala, Sweden.; Natural History Museum, University of Oslo, P.O. Box 1172, Blindern, 0318 Oslo, Norway.

## Abstract

The European Reference Genome Atlas (ERGA) consortium aims to generate a reference genome catalogue for all of Europe’s eukaryotic biodiversity. The biological material underlying this mission, the specimens and their derived samples, are provided through ERGA’s pan-European network. To demonstrate the community’s capability and capacity to realise ERGA’s ambitious mission, the ERGA Pilot project was initiated. In support of the ERGA Pilot effort to generate reference genomes for European biodiversity, the ERGA Sampling and Sample Processing committee (SSP) was formed by volunteer experts from ERGA’s member base. SSP aims to aid participating researchers through i) establishing standards for and collecting of sample/ specimen metadata; ii) prioritisation of species for genome sequencing; and iii) development of taxon-specific collection guidelines including logistics support. SSP serves as the entry point for sample providers to the ERGA genomic resource production infrastructure and guarantees that ERGA’s high-quality standards are upheld throughout sample collection and processing. With the volume of researchers, projects, consortia, and organisations with interests in genomics resources expanding, this manuscript shares important experiences and lessons learned during the development of standardised operational procedures and sample provider support. The manuscript details our experiences in incorporating the FAIR and CARE principles, species prioritisation, and workflow development, which could be useful to individuals as well as other initiatives.

## I. The Sampling and Sample Processing committee of ERGA

The European Reference Genome Atlas (<u>ERGA, Mazzoni et al. 2023</u>) consortium, the European node of the <u>Earth BioGenome Project</u> (EBP; Lewin et al. 2022), aims to generate a publicly available reference genome catalogue for all European eukaryotic biodiversity (Formenti et al. 2022; Theissinger et al. 2023). ERGA has the potential to catapult the fields of biodiversity conservation, evolution, ecology, and others to a new sphere analogous to how the first complete sequence of the human genome surged the fields of medical genetics, genomics, anthropology, and others (Formenti et al. 2022; Theissinger et al. 2023). It is akin to the appearance of the first natural history collections dating back as far as the 1800s that still lay the foundations for many new and important insights today.

ERGA is led by its chair and two co-chairs in cooperation with the ERGA council (a team consisting of two elected representatives of each member country). To support the multitude of ERGA tasks, <u>several scientific and Science+ committees</u> have been established. ERGA’s first project - <u>the ERGA Pilot (McCartney et al. 2023)</u>, tested a distributed genomics infrastructure while fuelling the ERGA committees. The Pilot Project is a community effort without a dedicated funding source, which will result in the production of over 98 genomes from 34 provider countries, connecting close to 400 involved ERGA members.

The Sampling and Sample Processing committee (<u>SSP</u>) is a committee of volunteer expert ERGA members tasked with developing guidelines to support sampling and sample processing. Specifically, the SSP’s initial responsibilities included i) establishing standards and mechanisms to collect sample/specimen metadata; ii) prioritising species collection; and iii) developing taxon-specific collection guidelines for the biological material underlying ERGA’s mission. The specimens and their derived samples are provided through ERGA’s large network of biodiversity partners spread across Europe (Box 1).

### Box1.

The scheme shows the ERGA workflow in the Pilot project. Species were initially nominated by the ERGA community (1), accompanied by a comprehensive form containing questions used for Species Selection (2), based on several exclusion, prioritisation and feasibility criteria. Species were distributed to the participating Sequencing Partners (3), which were responsible to contact the Genome Team lead (often the sample provider) to organise all necessary onboarding and regulatory requirements and documentation and agreed to generate reference genomes that fulfil EBP quality metrics (4). Samples were collected, vouchered, and several tubes of subsamples were prepared for sequencing as arranged with the sequencing partner and collaborating research groups (5). Sample providers were also encouraged to barcode the samples prior to sequencing and to store corresponding material in local biobanking facilities. Metadata was recorded using the ERGA sample manifest following established guidelines (6), uploaded to the metadata brokering platform COPO and validated by the Pilot sample management team (7). After confirmation that all the required documentation and metadata was in place, samples were shipped assuring a cold chain to the designated sequencing facility (8).

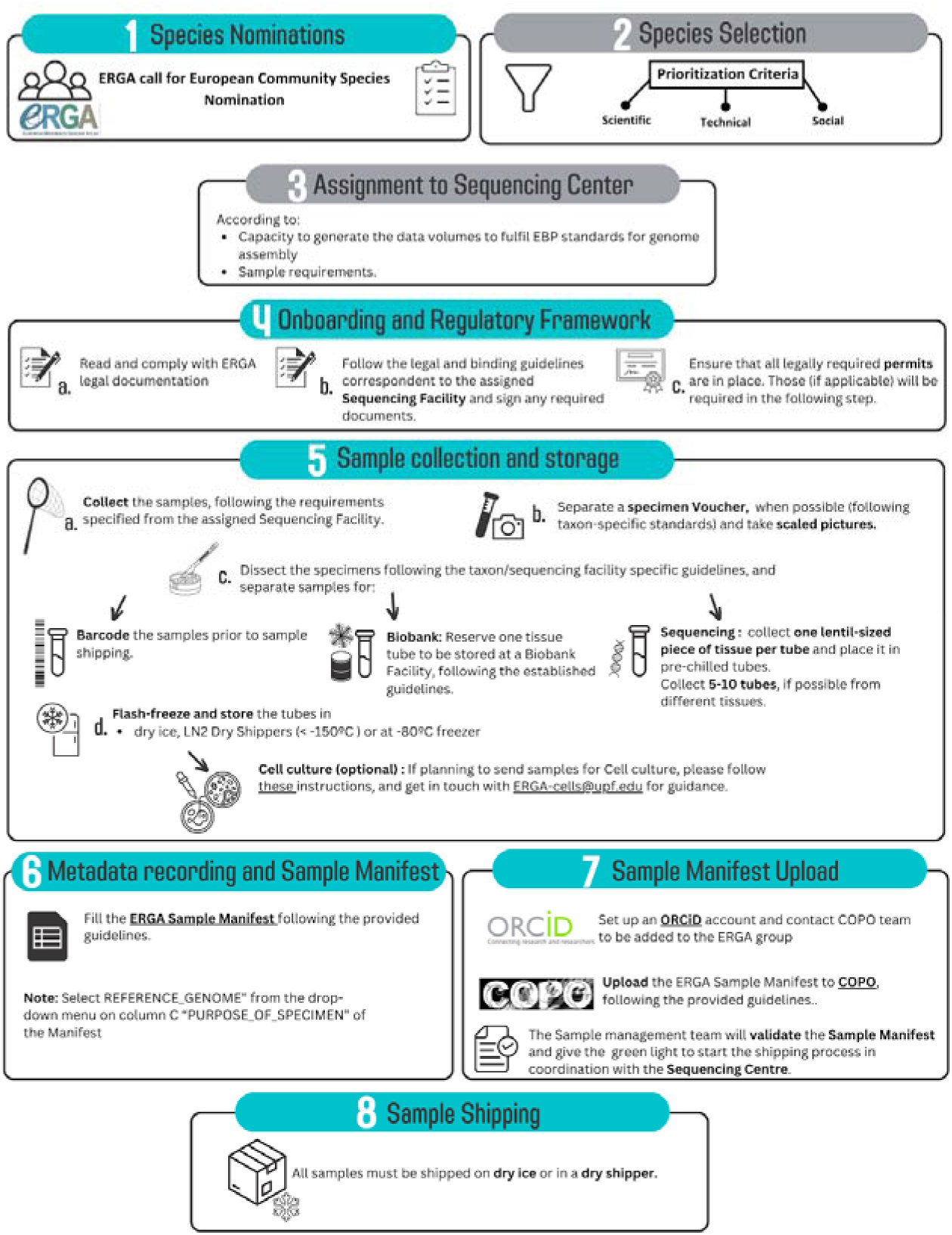

The SSP serves as the sample provider’s entry point into ERGA’s distributed genomic infrastructure and helps ensure standardised sample processing. As ERGA was maturing, additional SSP tasks emerged: iv) providing guidance to sample providers for the compliance with legal obligations in collaboration with ERGA’s <u>ELSI committee</u> (Ethical, Legal, and Social Issues) and v) sample provision - facilitating sample shipping between sample providers and sequencing centres.

As the number of EBP-associated projects across the globe gradually increases, we share here the experiences we gained whilst developing the operational procedures and sample provider support systems for the first continent-wide, distributed, genomics infrastructure. We hope our lessons can be useful to other large consortia who are pursuing the shared mission of sequencing all of life. Our experience in tackling <u>FAIR</u> (Findable, Accessible, Interoperable, Reusable) and <u>CARE</u> (Collective benefit, Authority to control, Responsibility, Ethics) data principles, species prioritisation, and workflow development may also be of use to smaller initiatives.

## II. The sample flow within ERGA

Reference genome production within a multinational consortium like ERGA involves many partners spanning dozens of countries. To manage diverse expectations, ensure efficient task execution, streamline communication, and safeguard fair attribution, ERGA has implemented the formation of multidisciplinary ‘Genome Teams’ (Supplementary File 1). These include all contributors to the production of a reference genome (i.e., researchers, stakeholders, and rights holders) from the field to the final data analysis. The Genome Team lead’s (in the ERGA Pilot known as the sample ambassador) initial responsibilities include providing all necessary documentation, data, and metadata for a sample to enter the sequencing workflow (Box 1). Most often, this function is filled by the sample provider. All members of the Genome Team agree to adhere to <u>ERGA’s Sample Code of Practice</u> as well as <u>ERGA’s Code of Conduct</u>. The SSP committee serves as an important touch point for the Genome Team lead, providing advice and guidance on sampling requirements, metadata standards, legal compliance, and vouchering strategies.

### Selecting species for biodiversity genomics - species prioritisation in ERGA’s initial phase

Reference genome sequencing initiatives require implementing prioritisation criteria, given resource and technical limitations that prevent sequencing all targeted species immediately. Scientific, technical, and social criteria can govern such species prioritisation.

**Table 1.**
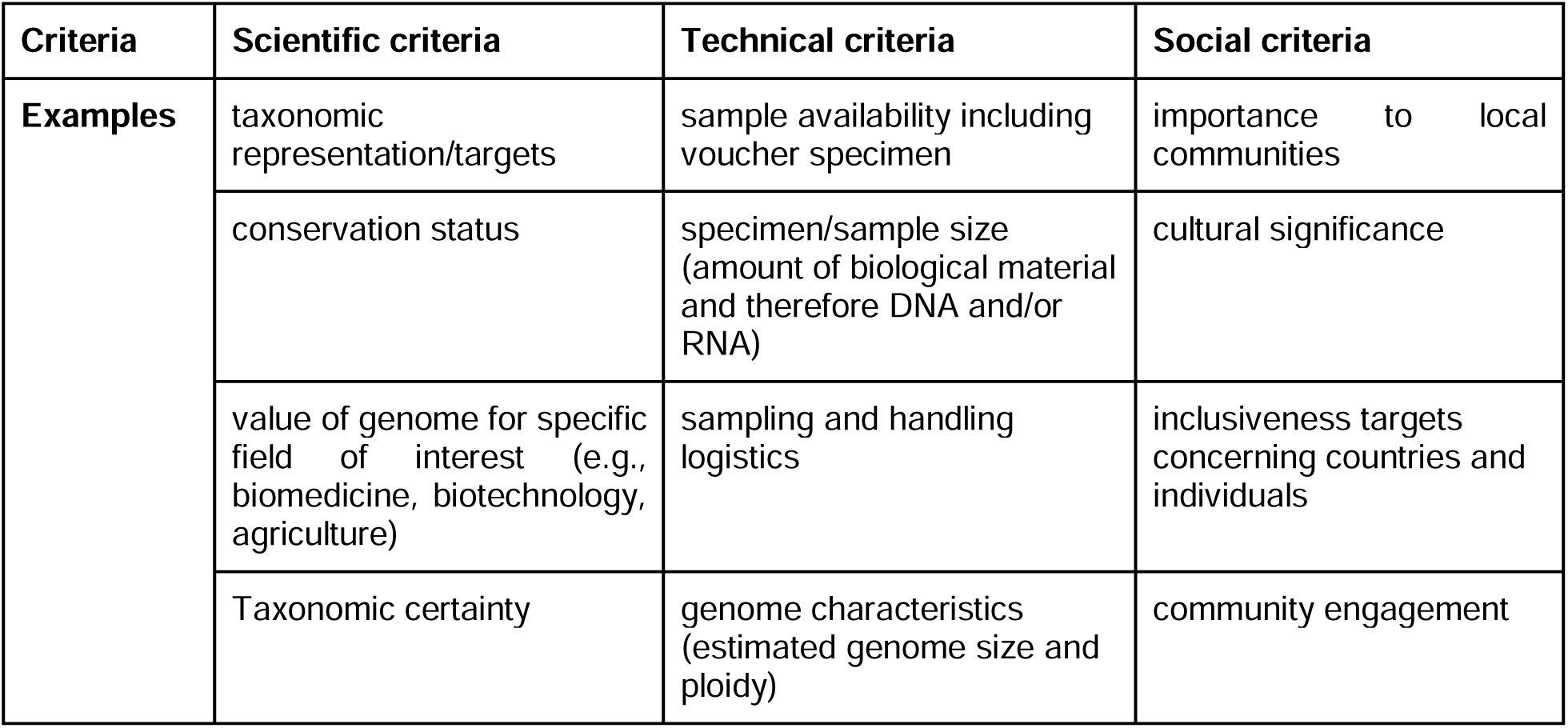
Non-exhaustive list of criteria for species prioritisation for genome sequencing projects.

| Criteria | Scientific criteria | Technical criteria | Social criteria |
| --- | --- | --- | --- |
| Examples | taxonomic representation/targets | sample availability including voucher specimen | importance to local communities |
|  | conservation status | specimen/sample size (amount of biological material and therefore DNA and/or RNA) | cultural significance |
|  | value of genome for specific field of interest (e.g., biomedicine, biotechnology, agriculture) | sampling and handling logistics | inclusiveness targets concerning countries and individuals |
|  | Taxonomic certainty | genome characteristics (estimated genome size and ploidy) | community engagement |

For initiating ERGA as a continent-wide genomic infrastructure network, a pool of candidate species for reference genome generation was solicited that were representative of the diversity of species and scientists across the consortium. To this aim, the ERGA community was asked to propose species through an initial simple ERGA species suggestion form resulting in 276 nominations. Subsequently, nominating persons were contacted to complete a comprehensive form (Supplementary File 2) containing 117 questions and commenting fields. The form included questions related to taxonomic identity, genome properties, voucher availability, habitat of species in question, sampling strategy, species conservation status, permits to obtain material for genome sequencing, sample properties (e.g., sex, amount, preservation quality, and tissue type), and species identification certainty. The refined species nomination form was open for 26 days and received 155 submissions.

After excluding species that already had available reference genomes, SSP implemented a prioritisation process based on country of origin and a simple scoring system, attributing a score of 1 to 3 in eight categories (Table 2). Higher priority was given to species that: i) had a genome size smaller than 1Gb, ii) were readily available, iii) could be freshly collected and for which biological material could be flash frozen, iv) could deliver >1g of tissue (if the organism permitted) and had well-established extraction protocols that allowed isolating chemically pure HMW DNA, v) could deposit a specimen voucher, vi) had no ambiguity risk in species identification, vii) had all permits present or were not needed (a formal documentation for either of the solutions was requested), and viii) had no export restrictions (if applicable).

**Table 2.** Feasibility criteria scoring for species suggested as sequencing targets of the ERGA Pilot Project.

| Category | 1 | 2 | 3 |
| --- | --- | --- | --- |
| **Genome size** | <1Gb | 1-3Gb | >3Gb |
| **Sample Availability** | Until end April 2020 | May-June 2020 | July 2020 or after |
| **Sample Preservation** | Freshly collected, flash frozen, -80°C, no preservative, never thawed | in-between 1 and 3 (to be evaluated by sequencing centre) | Not freshly collected and/or thawed several times, and/or not kept in -80°C |
| **Sample Size** | >1g | 100mg-1g | <100mg |
| **Suitability for HMW DNA** | Already extracted or taxon known to work well (e.g., vertebrates) | Not tested and not known for the taxon (can be checked with sequencing centres) | Inhibitors known to make DNA extraction and/or sequencing very challenging |
| **Voucher & SpeciesID** | Voucher kept in collection and no ambiguity in species identification |  | No voucher and/or ambiguous species identification |
| **Sampling Permits** | Yes or Not needed (documentation required either way) | Pending | No when needed or No documentation |
| **Export Regulations** | No restrictions between countries where sample will be handled or entire sequencing performed within country | Indexed to conservation status or Nagoya regulations to be clarified | No possibility for obtaining needed permits |

After ranking the species according to this scoring system, each proposing country was given the opportunity to refine their selection of species and to propose three final species considering three predefined target categories (endangered/iconic, marine/freshwater and pollinator) to match the available resources. At that point, ERGA had no centralised funding so feasibility was strongly determined by the availability of sufficient funds to support genome sequencing for a particular species. The project relied on resources contributed by participating ERGA members, institutions, and sequencing centres, with some additional support from industrial sponsors, that was used to supplement equity deserving genome teams in order to improve wide access to participation. As an extension to the selected list, standalone species were also included under the ERGA umbrella if they were completely funded by independent resources.

The circulation of the list of nominated species within ERGA resulted in cross-country collaborations especially for species proposed by more than one country, fostering exchange and reducing costs and redundancies.

The species selection and prioritisation process resulted in 98 selected species (https://goat.genomehubs.org/projects/ERGA), from 15 phyla (Figure 1B) and 34 countries or regions. With six of the seven selection scores relating to feasibility (including legal), this was the most prominent criterion, while the other criteria (i.e., conservation status, scientific relevance, socioeconomic relevance, taxonomic gaps, and community engagement) played only an indirect role via the subjective selection by the ERGA council members. ERGA has planned to implement unbiased species selection procedures in the future to alleviate the dominance of feasibility as selection criterion (see section V below).

**Figure 1.**
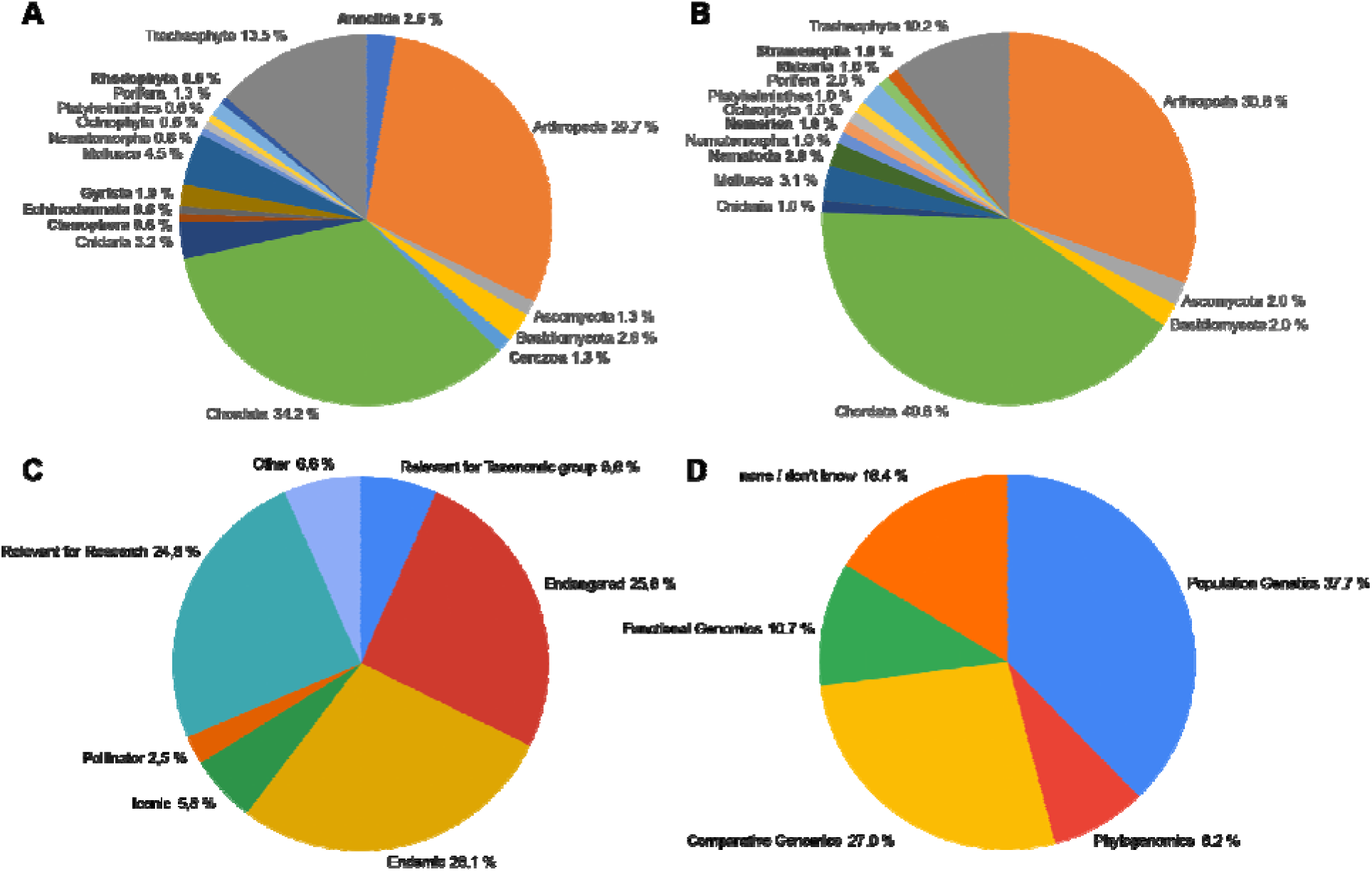
Pie charts of the number of species per phylum that were suggested for the ERGA Pilot Project at the beginning (A) and that are on the list of genomes realised or in production as of April 25th 2023 (B). The phyla are indicated together with the percentage of species per phylum. Phyla, which are different between A and B, are highlighted in bold. Additionally, the criterion for choosing the species (C) and the planned downstream analyses (D) are provided in percentages.

Both the initial and the final list of selected species showed a predominance of chordates, arthropods, and tracheophytes. Given that the initial pool of species was suggested by the ERGA community, this predominance may reflect the organism-bias of the biodiversity genomics community at large (see below). This taxon bias remained despite the dynamic nature of the taxonomic composition, as some species were removed due to sampling or sequencing technical barriers whilst others were added to increase representation and participation across ERGA’s diverse members. A total of 37% of the species were considered for the category endangered/iconic, and 12% were pollinators (as one example of scientific relevance and a target group of the Biodiversity Strategy of the European Commission). Most of the reference genomes were generated because the species are endemic (28%), endangered (26%) (and therefore the genome could be leveraged to inform conservation plans in the future) or to be used to answer specific scientific questions (25%) (Figure 1C). The most popular planned downstream analyses involve population genomics (38%) or comparative genomics (27%) (Figure 1D) (data from a questionnaire to species ambassadors, done by ERGA’s Data Analysis Committee, DAC, in the framework of Mc Cartney et al. (2023)).

Regarding inclusiveness, of the 18 <u>Widening countries</u> represented in the ERGA council 17 had at least one species included in the final list of generated reference genomes. The representation of ITC (Inclusiveness Target Countries) and Widening countries with 44 and 50 % of the 34 countries suggesting species is good overall. However, only 36 or 42 % of the final species came from ITC or Widening countries, respectively.

## II. FAIR and CARE principles, Metadata Collection and Brokering

### FAIR and CARE principles

As the number of initiatives working towards complete reference genomes for all of eukaryotic life are increasing, so too is the demand for freshly collected, wild specimens. This provides an opportune and pertinent moment to revisit biodiversity genomic metadata standards to ensure they are both scientifically comprehensive and also align with current ethical, legal and social standards for data governance. Ensuring that data are **f**indable, **a**ccessible, **i**nteroperable and **r**eusable (FAIR) is fast becoming a central dogma of the biodiversity genomics community (Wilkinson et al. 2016)^1^. Throughout the metadata standard development process (see next section), SSP intentionally and carefully aligned all ontologies to the FAIR principles to safeguard that all ERGA data would have a maximised scientific potential, increased re- usability, and greater longevity.

Indigenous Peoples and Indigenous knowledge systems have, and continue to be, treated as subordinate and outside of western science, specifically when considering contextual metadata (Turner 2022). This has had the systematic consequence of severing the connection between Indigenous Peoples and Local Communities with their samples and data. To mitigate the manifestation of this exclusion within ERGA, SSP developed new metadata ontologies to support the disclosure of Indigenous rights and interests by Indigenous Peoples by sample providers. This purposeful inclusion and recognition of Indigenous Peoples and their rights actualises the CARE principles of Indigenous data governance (Carroll et al. 2021) whilst simultaneously working in complementary fashion to the FAIR principles. By creating this space at the entry point into ERGA processes, i.e., sample provisioning, SSP provided an opportunity for Indigenous Peoples and knowledge systems to permeate throughout the process of reference genome production and beyond (Figure 2). By operationalizing the FAIR and CARE principles across the metadata ontologies developed, ERGA members are supported to responsibly and openly share data.

**Figure 2.**
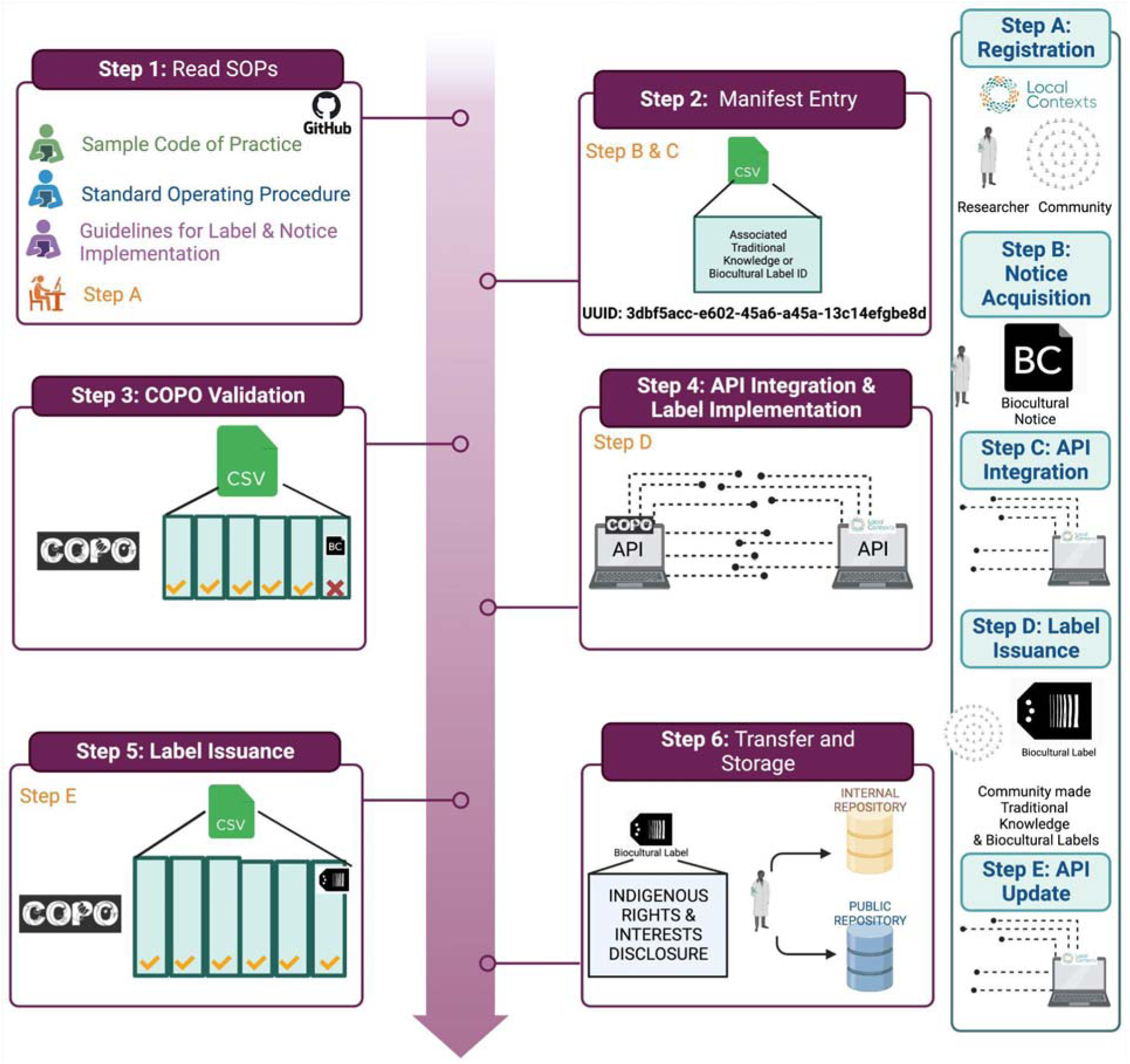
ERGA’s Biocultural and Traditional Knowledge Labels and Notices implementation protocol.

### ERGA Manifest for Metadata Collection and Brokering

Developing consortium-wide procedures for metadata collection is an opportunity to set a minimum standard of excellence, and ensures consistency across datasets. This approach is also a challenge since an unintentional exclusion of an important metric will lead to its systematic erasure from all data produced by the consortium. To support ERGA’s sampling process, SSP implemented the consortium’s first metadata standard, the <u>ERGA manifest</u>, and its supporting documentation (standard operating procedure (SOP)). This SOP and manifest were built on pre-existing standards that were developed for an established reference genome production initiative, <u>Darwin Tree of Life</u> (Lawniczak et al., 2022; Shaw et al., 2022), which followed the <u>Darwin Core standard</u>. The manifest supports ERGA’s goal to collect standardised, high-quality metadata that remains linked to the genome across the relevant repositories. The highly detailed SOP facilitates completing the ERGA manifest by the Genome Team lead who is responsible to provide information on: 1) sample identifiers (e.g., field and tube numbers, Genome Team lead), 2) taxonomic details, 3) sample type (e.g., life stage, organism part), 4) the sequencing partner, 5) sample collection event, 6) taxonomic identification and uncertainty, 7) sample preservation, 8) DNA barcoding, 9) biobanking and vouchering, 10) regulatory compliances including Indigenous rights and traditional knowledge, and 11) other relevant comments from the Genome Team representative.

The SOP explains every data point asked for, links to explanatory resources such as tutorial videos, and help contacts.

Expert members of SSP, i.e., sample managers, help genome teams upon request with filling in metadata fields and choosing appropriate terms in case of doubt. Sample managers can also check manifests prior to submission to avoid frustrating periods of trial and error for sample providers. Based on continuous user feedback, the SOP is updated twice a year to facilitate metadata collection for genome teams.

Upon upload of the manifest through the metadata brokering platform <u>COPO</u> (Shaw et al., 2020), metadata fields are validated against predefined standards and checklists to ensure terms and formats meet both ERGA and data repository expectations. Guidance to this process is provided through a visual guide on the COPO help webpage.

Upon manifest validation by the sample managers, an indicated set of mandatory metadata fields are brokered to the European Nucleotide Archive (ENA) under a dedicated BioSample entry ultimately connecting the digital sequence data to standardised sample metadata.

To mitigate the risk of missing information important to specific taxonomic groups or habitats due to own bias (see below), SSP included diverse team members when developing the manifest and planned for bi-annual updates of the metadata protocol so that accidental exclusions could be fixed in a timely manner and allow sufficient implementation and testing time for front- and backend development. Any issues with the manifest encountered by the community can be raised in the ERGA manifest GitHub or by contacting the SSP directly. The ERGA Pilot allowed the SSP committee to test the ERGA manifest on a broad variety of organisms by a pan-European network of researchers. Guidance for understanding and implementing the collection of metadata and vouchers was extensively requested and provided by SSP members. Finalisation of the ERGA manifest and its SOP was achieved through discussions with other ERGA committees, especially ELSI, and the ERGA coordination. The ERGA metadata collection is a semi-automated process that is highly scalable, preparing ERGA for an anticipated increased sample workflow. Validation of the sample manifest is the checkpoint of transitioning to the sequencing workflow.

The SSP data collection process links biological material, metadata, and sequence information in a maximally automatised fashion over open access databases and throughout the genome workflow from collection through nucleic acid extraction, sequencing, assembly and annotation steps. While open access genomic information is already a highly appreciated resource, comprehensive metadata enhances its value by making it more reusable. It is crucial that the metadata, sample(s), and derived sequence data are linked from the outset, because the opportunity to link them declines substantially with time (Crandall et al. 2022).

### Status Quo of metadata collection amongst biodiversity initiatives

To gain an understanding of the diversity and interoperability between the various metadata collection procedures being implemented within the community, SSP conducted a survey across global biodiversity genomics projects (Figure 3). A total of 24 initiatives that are actively generating high-quality reference genomes for non-human species responded. spanning Africa, North America, Oceania, Europe and Asia^2^*.

**Figure 3.**
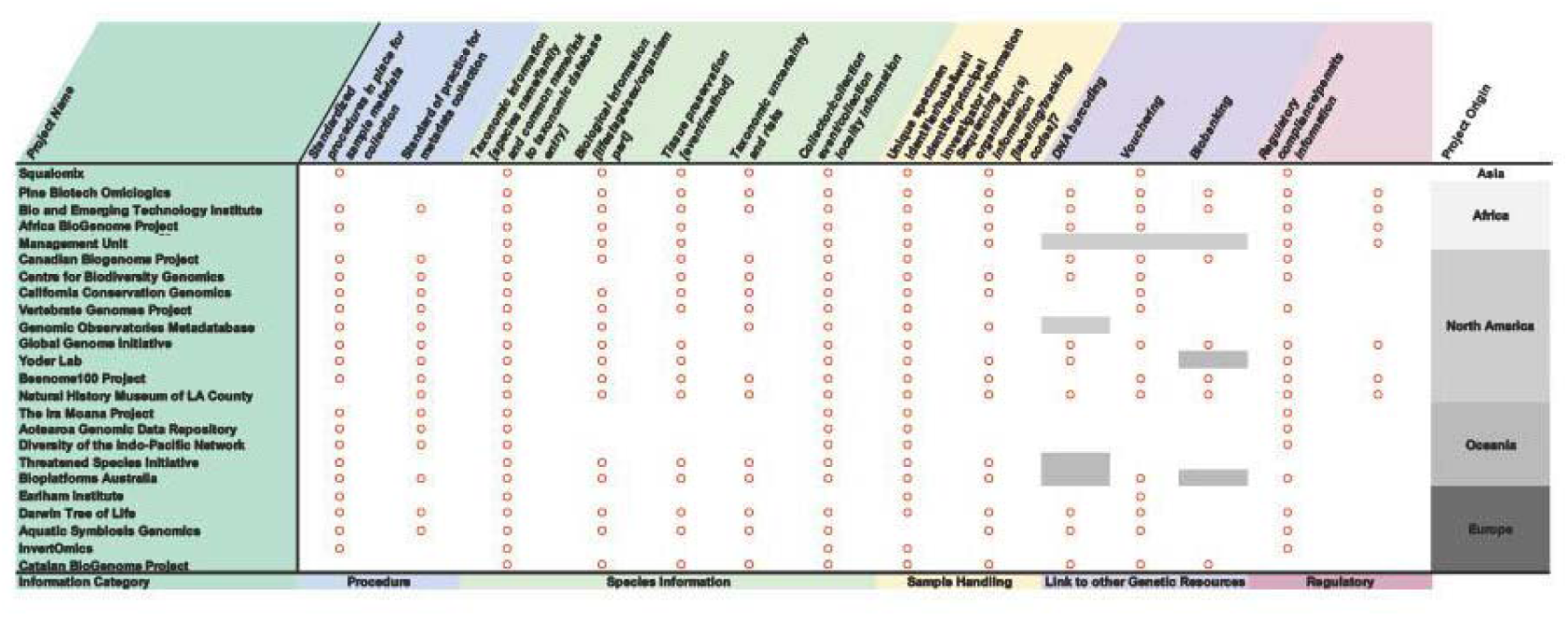
Results summary from the metadata survey conducted across 24 biodiversity initiatives worldwide. Red circles within a cell indicate presence, and empty cells indicate absence.

The results indicate that overall, 83% of responding initiatives have a standardised metadata collection procedure in place and 67% have an associated SOP to support and guide researchers in the metadata submission process. In terms of species-specific metadata collection, initiatives prioritise the collection of taxonomic (100%), collection information (96%), biological information (75%) and tissue preservation (75%) over providing more fine-grained information on the taxonomic uncertainty or risks associated with the species being sampled (59%). Almost all initiatives (96%) collected unique specimen and tube/well identifiers as well as the associated principal investigators whereas just 67% required information about the sequencing facility.

The amount of metadata collected about other associated genetic resources from the species sample was relatively low. For instance, only 55% of the 20 projects collect DNA barcoding information within their metadata. Further, just 65% of initiatives collect vouchers and 33% collect cryopreserved samples and require this information as part of their standard metadata collection processes. Finally, 42% of initiatives required some kind of disclosure of regulatory compliance and just 33% of projects required metadata concerning associated Indigenous rights and interest.

### Scaling Legal Compliance

SSP also focussed on creating an infrastructure that supports and promotes legal as well as ethical and scientifically sound sample collection. As an initial safeguard, SSP supported ERGA to develop a document of best practices for ethical and legal sample collection (<u>ERGA Code of Conduct</u>). All researchers participating in the Pilot were required to agree to these practices in advance of making their metadata manifest submission. These practices detailed expectations surrounding local, regional, national, and international permitting in addition to how to ethically collect samples to minimise harm.

Further, the ERGA manifest contained seven metadata fields regarding the regulation and permit requirements for each sample. These questions comprise comprehensively all permit forms that could be required to obtain a sample for genome sequencing: i) initial question if regulatory compliance is required and adhered to, ii) Applicability of traditional knowledge or biocultural rights with subsequent collection of rights definition, project ID provided by the Local Context Hub and contact information iii) Request for ethics permit applicability, definition and permit iv) Request for sampling permit applicability, definition and permit and v) Request for Nagoya Protocol permit applicability, definition and permit. This comprehensive request for applicability and documentation of compliance raises awareness also for sample providers to respect all regulations.

In partnership with COPO, ERGA required the mandatory upload of permits during the manifest submission process. Expert personnel within ERGA were alerted when a permit had been uploaded into the directory and, where possible, confirmed the appropriate permits had been obtained.

### The importance of vouchers for biodiversity genomics

Voucher specimens in natural history collections are benchmarks against which we compare the world around us. They illuminate how the world has been changing, and especially how we have been changing the world. Reference genomes are a new benchmark. Vouchering is critical to genomics because it provides a permanent, verifiable, and accessioned record of the identity of the organism being sequenced and, in some cases, a sample of its genetic material (biobanking). When determining which of the many available vouchering methods is most appropriate, consideration should be given to e.g., the taxon, its size, its conservation status (Table 3). The SSP determined that a sample voucher helps contextualise the biology of the organism and thus increases the probability that the sequencing data generated will be aligned with FAIR principles and useful into perpetuity.

**Table 3.**
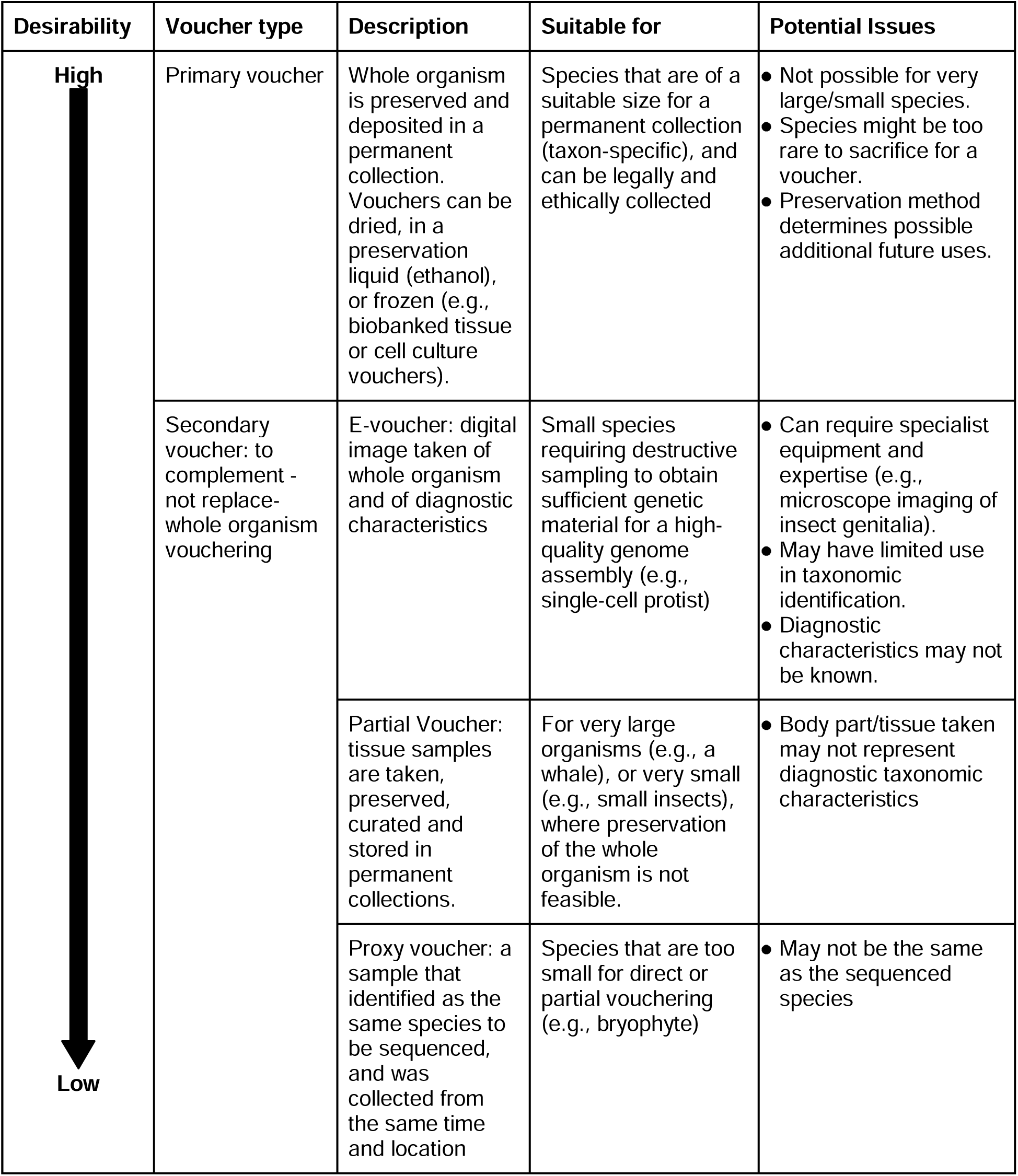
Vouchering methods available to specimens destined for genome sequencing. Note that multiple voucher types may be made for a single genome.

A driving rationale for vouchering is the fluid nature of taxonomy, as new scientific insights lead to changes in the classification of species. As this happens, the prescribed identity assigned to a sequenced individual could be questioned. In such cases, the presence of a voucher can be used to re-examine the species to confirm, or alternatively revise and update, its identity. Furthermore, vouchers can improve data quality assurance, reduce the risk of data corruption, and eliminate the propagation of confusion when a taxonomic revision has taken place.

Even for taxonomically stable groups, a voucher specimen provides the possibility to join morphological and genome sequence information and verifies the specimen/ species from which the genome was produced. A physical voucher can also be used for other analyses, including photographic, x-ray, CT imaging, and/or chemical analyses such as stable isotopes. A biobanked sample could unlock opportunities for future exploration (e.g., RNA, secondary genetic marker analyses such as methylation).

## IV. Sample provision: connecting genome teams with sequencing centres

Sampling and sample transfer can be a complicated endeavour with its multilayer complexity arising from four main categories: biological, logistic, administrative/policy and legal issues. These challenges can strongly influence the outcome of the project and impede the proper transfer of the samples to a sequencing centre (Box 1). The role of SSP is key to overcoming these issues and ensuring the legal, ethical, and timely flow of samples from sample collectors to sequencing centres (Figure 4).

**Figure 4.**
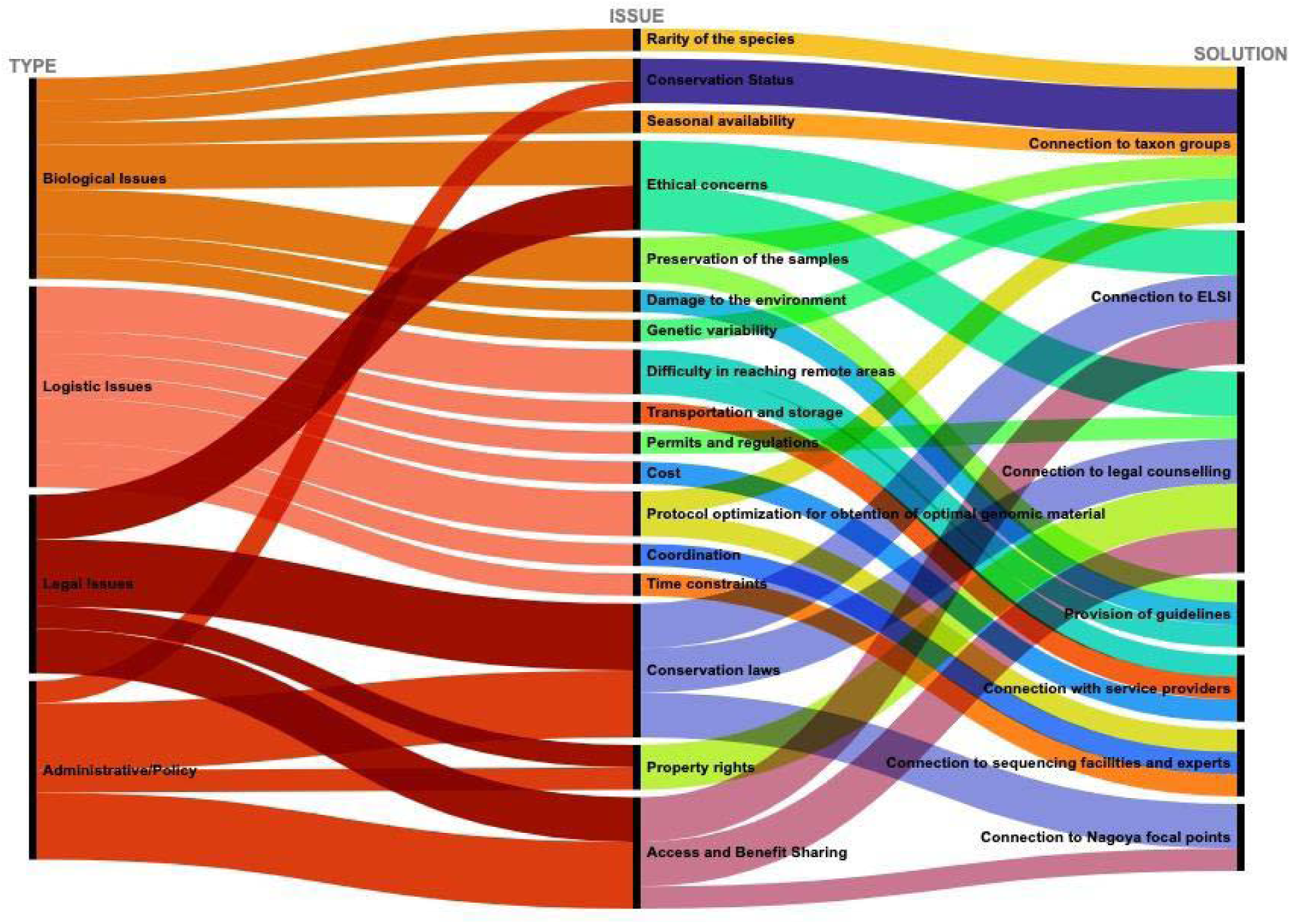
The role of SSP supporting critical issues prior to and after sample collection. Type of issue affecting sample provision, description of issues and solutions are indicated.

The distributed genomic infrastructure developed by ERGA promoted and supported the decentralisation of sequencing efforts across Europe. While many sampled species were sequenced within their country of origin, others were shipped to an international sequencing centre. Regardless of the length and duration of shipment involved, ERGA recommended cold-chain shipment, which is necessary to preserve the integrity of nucleic acids. Since this can be a challenge for sample providers, ERGA tried to connect sample providers with sequencing centres that were geographically close and aided in sample transportation within the ERGA network. Maintaining the integrity of nucleic acids is a prerequisite to meet the EBP standards of genome assembly utilising the current sequencing technology (Dahn et al. 2022). However, samples are often collected in remote locations, where access to appropriate courier service is financially not feasible or simply not available, a challenge that the ERGA Pilot also faced. Further, there is a series of legal procedures that require consideration to ensure compliance with regulations and safety standards, including, among others, chain of custody forms (to document the movement of the samples from collection to sequencing), material transfer agreements (a legal contract between two parties that governs the physical transfer of the biological samples between them, and which establishes the terms and conditions under which the materials will be transferred), import/ export permits (that may be required depending on the country of origin and destination), health certificates (required by some countries to ensure that the samples do not pose a risk to human or animal health), and/or CITES permits (required if the samples are from a species protected under the Convention on International Trade in Endangered Species of Wild Fauna and Flora), as well as ABS/ Nagoya relevant national implementations, among others. The ERGA Pilot project served as an opportunity to understand the magnitude and complexity of these needs and actions in a collective manner, with everyone implicated learning about pieces of information that could make an impact in the success of the full logistics chain. For instance, we learned that different shipping companies operate better in certain geographical regions, and that sometimes it is important to ask them explicitly to refill the dry ice during the transit. We also collectively learned about the bureaucratic idiosyncrasy of each country with respect to export and import permits and Nagoya protocol, with some countries being more flexible and others being more restrictive. All these pieces of information have been shared with SSP and are being leveraged to develop SOPs to facilitate the transit from species collectors to sequencing centres, and will have a strong impact in the implementation of larger projects such as Biodiversity Genomics Europe (see below).

### Future taxon-specific best-practice guidelines

The biological diversity being sampled by large genome initiatives like ERGA necessitates the development of targeted best-practice sampling guidelines. The approach of having different sampling procedures for different taxa is very commendable as it eliminates complications arising from structural and functional variations between the taxa.

Such guidelines are imperative to ensure that sampling efforts minimise the number of samples taken, maximise the data quality, and increase the scientific utility of the sample. To this end, the SSP will take a taxonomic approach that seeks to balance providing a set of guidelines that are comprehensive, with enough specificity to support fit-for-purpose sampling, while simultaneously not providing too much information and materials that may overwhelm field biologists.

To develop these guidelines, separate working groups have been set up for each of the following broad taxon groups: vascular plants, bryophytes and macroalgae, macroinvertebrates, protists, soft bodied invertebrates, fungi and lichens, chordates, and arthropods. The goal of each group is to create a working protocol for the sampling of specimens within that taxonomic group, and those will follow a set structure to ensure consistency and readability. There is a strong foundation for these protocols (e.g. <u>dx.doi.org/10.17504/protocols.io.261gennyog47/v1</u>). ERGA has the intention of publishing these guidelines in open access over protocols.io

A key challenge in developing these guidelines will be to identify and include experts - taxonomic, field, and wet lab biologists- who are willing to voluntarily contribute their time and knowledge to the wider community. The SSP has reached out to the ERGA repeatedly to gain insight into ERGA members’ expertise and connect those to SSP. Based on this effort, SSP establishes communication with sample providers and ERGA member institutions that can provide expertise in e.g. sample handling, storing and species identification. This help is provided over the SSP email contact as well as a dedicated channel in the communication platform keybase (https://keybase.io/team/erga.listserv). *Vice versa*, a future challenge will be to work towards an adoption of these guidelines by the biodiversity community at large. Integrating, documenting, and distributing this knowledge and ‘know-how’ is fundamental to ERGA and its umbrella organisation, the EBP. Based on experiences in the ERGA pilot, members of the SSP and the ERGA BGE project consult with the EBP samples committee and the EBP executive board in areas where ERGA sees a need for larger adoption of processes and standards.

## V. Critical Bias Assessment

The biodiversity genomics community is subject to systematic biases that affect the accuracy and completeness of the produced data, and may limit the meaningfulness of the conclusions obtained. Bias comes in many forms, which have different impacts. The ELSI/ JEDI committee is more focused on the human dimension, and the SSP committee focused on country representation and taxonomic biases described here. ERGA as a consortium of European researchers is at its foundation intentionally geographically biased, while at the same time promoting and extending representation and participation of researchers across Europe. In the Pilot, prioritising this aim over the taxonomic breadth of the generated reference genomes resulted in the manifestation of taxonomic biases (see above).

Unbalanced representation of genomes being sequenced across the tree of life is common in biodiversity genomics initiatives, causing over-representation of some taxa with data available in public repositories. Non-model organisms and more “difficult” samples remain under-investigated because there are few standardised sampling collection, preservation, HMW-DNA extraction, and library preparation protocols available to manage non-optimal situations (e.g., small size, existence of exoskeleton or spicules, presence of substances that impair adequate DNA extraction or sequencing, etc.). This lack of knowledge on certain taxa reflects the available taxonomic expertise. For example, experts in vertebrates, certain arthropod and plant groups are vastly more abundant than for other large taxonomic groups like mollusks, nematodes or annelids (Capa & Hutchings 2021; Engel et al. 2021), which SSP quickly realised while forming taxon expert groups (see above).

Beyond taxonomy, other sources of representation bias exist in reference genome projects. Sample bias can occur when samples do not accurately represent the known or unknown heterogeneity of the taxon being studied. SSP encourages sampling from the type locality. Habitat bias occurs when samples are more often collected in certain types of habitats that are more common or more easily accessible, under-representing knowledge about habitat-specific species (e.g., caves, deep-sea). ERGA aims to target this bias with <u>calls for funded field expeditions</u> to understudied hotspots of biodiversity in Europe. Historical bias can have strong impacts, as samples collected based on prior knowledge or historical information may not accurately reflect the current state of diversity.

A prime goal of SSP is to raise awareness of the importance of taxonomic representation for genomics, and biodiversity research more generally, and the study of research deserving groups, species, populations and habitats. SSP has played a key role in creating a bridge between taxonomy- and taxon-specific experts with sequencing centres, and aims to create the conditions to explore the feasibility of genome sequencing for all eukaryotes. Biodiversity genomics benefits the most when it is inclusive in all aspects. Many hotspots of biodiversity exist in Europe, and many are positioned in nations and regions that are deserving of additional support. By creating a European-wide network, SSP aims to support such regions through capacity and capability building for genomics.

## VI. Where do we head?

We believe that overall, sequencing and assembling the initial cohort of species that entered into ERGA’s process was a success story. To a large extent this is thanks to collaboration and alignment with preexisting, well established biodiversity consortia e.g., DToL. Similarly, we hope that our prioritisation efforts, the ERGA metadata manifest, as well as the stewardship of legal, FAIR and CARE information, can be utilised, improved, or adopted by other biodiversity genomics projects, national or international, irrespective of the project size. An immediate example of this is the EU-funded project <u>BGE - Biodiversity Genomics Europe</u>, for which the ERGA initial phase has set the ground for key procedures of the sampling and sample processing process. The BGE consortium unites ERGA with the DNA barcoding community (<u>BIOSCAN Europe</u>) to promote the use of genomics to study and monitor biodiversity and create tools to tackle its decline. BGE will establish ERGA as the European node of the <u>Earth Biogenome Project</u> and formalise coordinated efforts, infrastructures and workflows to generate reference genomes of European species.

### Towards a balanced and strategic prioritisation of species

As ERGA moves forward, the biases identified are being reflected upon to iteratively improve sampling and prioritisation. As dedicated projects are established, such as BGE, the selection and prioritisation of species for reference genome generation can better approximate governing principles (see above “Selecting species for biodiversity genomics projects”), and be less dependent on circumstantial feasibility aspects and funding availability for particular taxa. These governing principles can be explicitly and objectively included into the species prioritisation process and with a more prominent role, while feasibility will likely remain an important aspect of species selection. Once priorities are established and weighted, the species selection process can be fully automated. Building on the first experiences of ERGA, such a process is being implemented in BGE. This process, which is developed with the larger ERGA community, gives more weight to taxonomic diversity, country of sample origin, countries with little representation in ERGA and involves sample providers using JEDI criteria (favouring novel sample providers, underrepresented groups, and involvement of non-scientific communities) and applicability of produced genome resource, followed by a check for technical feasibility. ERGA is displaying its target species over the platform Genomes on a Tree (https://goat.genomehubs.org/projects/ERGA), in agreement with other nodes of the EBP. ERGA members as well as SSP sample managers engage with other genome initiatives when overlaps are detected and facilitate collaboration in order to prevent parallel efforts.

### A live and comprehensive sampling metadata manifest

The ERGA metadata manifest and its SOP are living documents, which are regularly revised under strict version control (https://github.com/ERGA-consortium/ERGA-sample-manifest). During the Pilot phase, it became clear that the metadata core was not entirely comprehensive. For example, the first version could not capture sampling depth and only allowed inputting a precise location. This information is important in the marine context as it was not possible to correctly represent samples from trawls or transects. Updated releases of the manifest have acknowledged these gaps and now comprise fields for e.g., depth and latitudinal and longitudinal coordinates for two points instead of one for sampling transects, extended vocabulary for sampled tissues, etc. As ERGA progresses, adding more extensions might be necessary during the planned regular updates.

The question that is often raised in regard to metadata collection is what is the trade-off between comprehensiveness *versus* feasibility. Sampling for reference genome generation has many logistical steps that are important to document in the metadata record. Such extensive collection of metadata appears doable when the emphasis is on single (or a few) representative samples per species while we acknowledge that feasibility and applicability might be different for e.g., population data or already collected material that cannot be obtained again. Yet, as the field of genomics moves forward and technological advances allow extracting more data at higher quality from material with varying quality samples, extending the high ERGA standards to any sample collected for genetic analyses appears as an appropriate perspective. In this light, the increase in frozen archives that ERGA supports will be a treasure trove for genome initiatives.

### Streamlining legal compliance procedures

Biodiversity knows no boundaries and it is blissfully unaware of its traversal distribution across many national, political, and cultural borders that may have varying legal systems. However, ERGA is obligated to respect these borders and the legal systems within, and so a harmonisation of procedures will be a crucial aspect of building a streamlined European sampling infrastructure for reference genome generation. ERGA’s network provides cross-country communication, which should be extended to local authorities, and ensure efficient flow of information about specific legal requirements of sampling. Streamlining the steps required to ensure legal compliance therefore is an important way to increase the efficiency of the reference genome generation pipeline.

### A continued concerted effort

Under the umbrella of the EBP and in the light of the progress that sequencing technology and data processing offer, there is a need to scale up the genome generation process. While ERGA has pioneered the establishment of a collaborative transnational effort for reference genome generation in Europe, other regional initiatives advance and face similar challenges. We here call for the establishment of collaborative concerted efforts among different consortia under the EBP flag, unifying standards across the whole workflow, starting with sampling and sampling processing and ending with making data available via open repositories.

## Acknowledgements

We thank all members of the ERGA SSP committee and the committee’s meeting participants for their support to the SSP and ERGA mission. In particular, we thank Alice Minotto and Felix Shaw from COPO, Josephine Burgin and Joana Pauperio from the EBI/EMBL, as well as Luisa Marins (Leibniz Institute for Zoo and Wildlife Research) for their help in implementing the ERGA manifest. We thank the Samples Working Group of the Darwin Tree of Life Project for a fruitful exchange on metadata collection and standards. We thank ERGA’s Data Analysis Committee for access to questionnaire data used in Figure 1. We acknowledge the essential work of Giulio Formenti and Alice Mouton, members of the ERGA Pilot Project coordination team, in making this work possible by contributing to build the necessary sample metadata collection infrastructure, including the early establishment of the sample manifest collection process, the ERGA manifest Github repository, as well as with their constant coordination effort as part of the ERGA Pilot Project daily activities. We especially thank the ERGA chairs for fruitful exchanges and their continuous support during the establishment phase of ERGA.

R. Oomen was supported by the James S. McDonnell Foundation 21st Century Postdoctoral Research Fellowship, the Natural Sciences and Engineering Research Council of Canada Postdoctoral Research Fellowship, and the Research Council of Norway (Earth BioGenome Project Norway; Project no. 326819). R. Fernández acknowledges support from the following sources of funding: Ramón y Cajal fellowship (grant agreement no. RYC-2017-22492 funded by MCIN/AEI /10.13039/501100011033 and ESF ‘Investing in your future’), project PID2019-108824GA-I00 funded by MCIN/AEI/10.13039/501100011033, and by the European Research Council (ERC) under the European’s Union’s Horizon 2020 research and innovation programme (grant agreement no. 948281). O. Vinnere Pettersson is supported by RFI/VR and Science for Life Laboratory, Sweden. S. McTaggart was supported by the Biotechnology and Biological Sciences Research Council (BBSRC), part of UK Research and Innovation, through the Core Capability Grant BB/CCG1720/1 and the Earlham Institute Strategic Programme Grant Decoding Biodiversity BBX011089/1 and BBS/E/ER/230002B. J. Melo-Ferreira acknowledges support from FCT, Fundação para a Ciência e a Tecnologia (2021.00150.CEECIND contract and project HybridChange, PTDC/BIA-EVL/1307/2020, via Portuguese national funds). T. H. Struck acknowledges funding from the Research Council Norway (project number 300587).

A. Böhne acknowledges support from the German Research Foundation DFG (grant numbers DFG 497674620 and DFG 492407022) and the Leibniz Association.

A. Böhne, R. Monteiro, R. Oomen, T. Struck, R. Fernandez, S. McTaggart, J. Melo-Ferreira, J. A. Leonard and O. Vinnere Pettersson were funded by Horizon Europe under the Biodiversity, Circular Economy and Environment (REA.B.3); co-funded by the Swiss State Secretariat for Education, Research and Innovation (SERI) under contract number 22.00173; and by the UK Research and Innovation (UKRI) under the Department for Business, Energy and Industrial Strategy’s Horizon Europe Guarantee Scheme. We would also like to acknowledge the contributions of the biodiversity genomics initiatives that contributed data concerning their metadata acquisition processes including Pine Biotech Omiclogics, Vertebrate Genomes Project, Bio and Emerging Technology Institute, The Ira Moana Project, Aotearoa Genomic Data Repository, Diversity of the Indo-Pacific Network, Earlham Institute, the Yoder Lab, Catalan BioGenome Project, African BioGenome Project, Squalomix University of Port Harcourt, Global Genome Initiative, Genomic Observatories Database, Beenome 100 Project, National History Museum of Los Angeles County, Threatened Species Initiative, Bioplatforms Australia, Darwin Tree of Life, Aquatic Symbiosis Project, and InverOmics.

Traditional Knowledge and Biocultural Label and Notice development: The implementation of the Labels and Notices and the development of the supporting guidance documentation was funded through the European Open Science Cloud (RDA_OSF_EOSC-228) in partnership with the Global Indigenous Data Alliance, RDA and Local Contexts.

## Glossary

| **Acronym** | **Explanation** | **Resource** |
| --- | --- | --- |
| ABS | Access and Benefit-Sharing | https://absch.cbd.int/ |
| BGE | Biodiversity Genomics Europe | https://biodiversitygenomics.eu/ |
| BIOSCAN EUROPE | part of the International Barcode of Life Consortium (iBOL) | https://www.bioscaneurope.org/ |
| CARE | Collective benefit, Authority to control, Responsibility and Ethics | https://www.gida-global.org/care |
| CITES | Convention on International Trade in Endangered Species of Wild Fauna and Flora | https://cites.org |
| COPO | Collaborative OPen Omics | https://copo-project.org/ |
| DToL | Darwin Tree of Life | https://www.darwintreeoflife.org/ |
| EBP | Earth Biogenome Project | https://www.earthbiogenome.org/ |
| DAC | Data Analysis Committee | https://www.erga-biodiversity.eu/team-1/dac---data-analysis-committee |
| ELSI | Ethical, Legal, and Social Issues | https://www.erga-biodiversity.eu/team-1/elsi---ethical%2C-legal%2C-and-social-issues |
| ENA | European Nucleotide Archive | https://www.ebi.ac.uk/ena/browser/home |
| ERGA | European Reference Genome Atlas | https://www.erga-biodiversity.eu/ |
| FAIR | Findable, Accessible, Interoperable, and Reusable | https://www.go-fair.org/fair-principles/ |
| GoaT | Genomes on a Tree | https://goat.genomehubs.org/ |
| ITC | Inclusiveness Target Countries | - |
| JEDI | Justice, Equity, Diversity & Inclusion | https://jedicollaborative.com/ |
| SOP | Standard Operating Procedure | - |
| SSP | Sampling & Sample Processing Committee | https://www.erga-biodiversity.eu/team-1/ssp---sampling-%26-sample-processing |

## Footnotes

1 FAIR was introduced by Wilkinson et al. (2016), which has since been accessed 580,000 times and cited 5,636 times

2 Notably, the lowest amounts of survey responses were obtained from Asia (the authors note that this is certainly due to our inability to identify appropriate contact points and does not reflect a lower number of biodiversity projects in this continent)

## References

Capa, M., & Hutchings, P. (2021). Systematics and Diversity of Annelids. MDPI, Basel. 10.3390/books978-3-0365-1389-8

Carroll, S. R., Herczog, E., Hudson, M., Russell, K., & Stall, S. (2021). Operationalizing the CARE and FAIR Principles for Indigenous data futures. Scientific Data, 8(1), Article 1. 10.1038/s41597-021-00892-0

Crandall, E. D., Toczydlowski, R. H., Liggins, L., Holmes, A. E., Ghoojaei, M., Gaither, M. R., Wham, B. E., Pritt, A. L., Noble, C., Anderson, T. J., Barton, R. L., Berg, J. T., Beskid, S. G., Delgado, A., Farrell, E., Himmelsbach, N., Queeno, S. R., Trinh, T., Weyand, C., … Toonen, R. J. (2023). Importance of timely metadata curation to the global surveillance of genetic diversity. Conservation Biology, 00, e14061. 10.1111/cobi.14061

Dahn, H. A., Mountcastle, J., Balacco, J., Winkler, S., Bista, I., Schmitt, A. D., Pettersson, O. V., Formenti, G., Oliver, K., Smith, M., Tan, W., Kraus, A., Mac, S., Komoroske, L. M., Lama, T., Crawford, A. J., Murphy, R. W., Brown, S., Scott, A. F., … Fedrigo, O. (2022). Benchmarking ultra-high molecular weight DNA preservation methods for long-read and long-range sequencing. GigaScience, 11, giac068. 10.1093/gigascience/giac068

Engel, M. S., Ceríaco, L. M. P., Daniel, G. M., Dellapé, P. M., Löbl, I., Marinov, M., Reis, R. E., Young, M. T., Dubois, A., Agarwal, I., Lehmann A., P., Alvarado, M., Alvarez, N., Andreone, F., Araujo-Vieira, K., Ascher, J. S., Baêta, D., Baldo, D., Bandeira, S. A., … Zacharie, C. K. (2021). The taxonomic impediment: A shortage of taxonomists, not the lack of technical approaches. Zoological Journal of the Linnean Society, 193(2), 381–387. 10.1093/zoolinnean/zlab072

Formenti, G., Theissinger, K., Fernandes, C., Bista, I., Bombarely, A., Bleidorn, C., Ciofi, C., Crottini, A., Godoy, J. A., Höglund, J., Malukiewicz, J., Mouton, A., Oomen, R. A., Paez, S., Palsbøll, P. J., Pampoulie, C., Ruiz-López, M. J., Svardal, H., Theofanopoulou, C., … Zammit, G. (2022). The era of reference genomes in conservation genomics. Trends in Ecology & Evolution, 37(3), 197–202. 10.1016/j.tree.2021.11.008

Lawniczak, M., Davey, R., Rajan, J., Pereira-da-Conceicoa, L., Kilias, E., Hollingsworth, P., Barnes, I., Allen, H., Blaxter, M., Burgin, J., Broad, G., Crowley, L., Gaya, E., Holroyd, N., Lewis, O., McTaggart, S., Mieszkowska, N., Minotto, A., Shaw, F., & Sivess, L. (2022). Specimen and sample metadata standards for biodiversity genomics: A proposal from the Darwin Tree of Life project. Wellcome Open Research, 7, 187. 10.12688/wellcomeopenres.17605.1

Lewin, H. A., Richards, S., Lieberman Aiden, E., Allende, M. L., Archibald, J. M., Bálint, M., Barker, K. B., Baumgartner, B., Belov, K., Bertorelle, G., Blaxter, M. L., Cai, J., Caperello, N. D., Carlson, K., Castilla-Rubio, J. C., Chaw, S.-M., Chen, L., Childers, A. K., Coddington, J. A., … Zhang, G. (2022). The Earth BioGenome Project 2020: Starting the clock. Proceedings of the National Academy of Sciences, 119(4), e2115635118. 10.1073/pnas.2115635118

Mazzoni, C. J., Ciofi, C., Waterhouse, R. M. et al. (2023). Biodiversity: an atlas of European reference genomes. Nature 619, 252. 10.1038/d41586-023-02229-w

McCartney, A. M., Formenti, G., Mouton, A. et al. (2023) The European Reference Genome Atlas: piloting a decentralised approach to equitable biodiversity genomics. bioRxiv 2023.09.25.559365. 10.1101/2023.09.25.559365

Shaw, F., Etuk, A., Minotto, A., González-Beltrán, A., Johnson, D., Rocca-Serra, P., Laporte, M.-A., Arnaud, E., Devare, M., Kersey, P., Sansone, S.-A., & Davey, R. (2020). COPO: A metadata platform for brokering FAIR data in the life sciences. F1000Research, 9, 495. 10.12688/f1000research.23889.1

Shaw, F., Minotto, A., McTaggart, S., Providence, A., Harrison, P., Pauperio, J., Rajan, J., Burgin, J., Cochrane, G., Kilias, E., Lawniczak, M., & Davey, R. (2022). Managing sample metadata for biodiversity: Considerations from the Darwin Tree of Life project. Wellcome Open Research, 7(279). 10.12688/wellcomeopenres.18499.1

Theissinger, K., Fernandes, C., Formenti, G., Bista, I., Berg, P. R., Bleidorn, C., Bombarely, A., Crottini, A., Gallo, G. R., Godoy, J. A., Jentoft, S., Malukiewicz, J., Mouton, A., Oomen, R. A., Paez, S., Palsbøll, P. J., Pampoulie, C., Ruiz-López, M. J., Secomandi, S., … Zammit, G. (2023). How genomics can help biodiversity conservation. Trends in Genetics, 39(7), 545–559. 10.1016/j.tig.2023.01.005

Turner, H. (2022). Cataloguing Culture: Legacies of Colonialism in Museum Documentation. University of British Columbia Press. https://press.uchicago.edu/ucp/books/book/distributed/C/bo70117236.html

