## Supplementary File 1 for "Contextualising samples: Supporting reference genomes of European biodiversity through sample and associated metadata collection"

### Supplementary File 1: Genome Team definition for the ERGA Pilot project

| Team Member | Role |
|---|---|
| Principal Investigator | Each genome has a designated Principal Investigator (PI) who is responsible to ensure the coordination of the project. If a sample ambassador specifically requests to lead the species genome analysis, this person will become the PI for this species' genome. Participants covering the costs for the genomes can also request to be PI or co-PIs, in which case the sample ambassador must be in agreement. In case the participants that are covering sequencing costs are not based in the same country as the sample origin, council members from the country of origin must agree to the request. Other participants can also request to become PI or co-PIs. The sample ambassador, in agreement with other potential co-PIs, decides on the request. Co-PIs appoint a coordinator among them responsible for the project, along with organizing the handling of the samples, to ensure that proper documentation meeting the Nagoya Protocol and local regulations is provided. |
| Sample collector and/or sample provider | Responsible for the ethical and legal collection of samples for ERGA and compliance with ERGA's 'Sample Collection Code of Best Practices' and 'Data Sharing and Management Policy'. |
| Taxonomist and/or ex-situ sample manager | Individual(s) responsible for ensuring the taxonomic validity of the sample obtained and its deposition into an appropriate biobank or collection understanding that the PUID associated with both will be disclosed in the ERGA metadata manifest. |
| Sample ambassador | The sample ambassador coordinates and organizes all of the samples, permits, barcoding, and storage components of the project, up to and including the shipment to laboratory and storage of vouchers. The individual (s) should be based in the country of origin or have an established research project in the area of sampling that justifies the involvement in genome establishment for species from another country. If not based in the country, s/he must provide proof of compliance with CBD Nagoya protocol as well as permits for in-country sampling, sample import/export and handling. The sample ambassador can also act as the sample provider. |
| Wet-lab processor | Facilitates all wet lab components of the project including HMW DNA extraction, library preps, sequencing. This is a hands-on researcher(s), or group, or could even be a facility PI. |
| Genome assembly manager | The individual(s) responsible that oversees the development, optimization, and correction of the genome assembly. Role also ensures appropriate computational resources are available. |
| Genome assembly and/or curation generator | The individual(s) responsible for the generation of the genome assembly (hands-on role). |

| | |
|---|---|
| Genome annotation and/or genome analysis generator | The individual(s) responsible for the generation of the gene annotation and data analysis (hands-on role). |
